# A novel enteric pacemaker activity in *Drosophila*

**DOI:** 10.64898/2025.12.04.690637

**Authors:** Niccolò P. Pampaloni, Isabella S. Balles, Paula Düllmann, Niclas Gimber, Eric Reynolds, Anatoli Ender, Jan Schmoranzer, Igor Iatsenko, David Owald

## Abstract

Enteric organs such as the gut and stomach display the intrinsic ability to generate slow-wave potentials, rhythmic electrochemical events acting as key regulators of gastrointestinal motility. In mammals, slow waves occur as a result of the pacemaker activity of Interstitial Cells of Cajal (ICCs) which, by recruiting visceral muscles, ultimately lead to peristalsis. Here, we uncover and characterize a novel mechanism in *Drosophila* by which the midgut is able to generate strong muscular whip-like contractions. This pacemaker activity, here named superstalsis, is only active at certain physiological conditions and is selectively linked to longitudinal visceral muscle activation. Furthermore, we identify in *Subdued*, an ortholog gene of the ANO1/TMEM16A family, a marker for peristalsis in *Drosophila*, specifically associated with circular muscle activity. As fasting and bacterial infections cause specific changes in the intestinal motility profiles *in vivo*, we propose that the gut can autonomously alter the peristalsis/superstalsis balance to efficiently respond to diverse pathological and homeostatic events.

## Main

Food ingestion and consequent nutrient absorption constitutes a vital mechanism for all animals. Heterotroph organisms, including mammals and flies, evolved complex specialized organs and mechanisms to finely regulate this process ^1,2^. As such, the gastrointestinal system displays specific autonomic visceral movements, a process called peristalsis, aimed to push forward and mechanically decompose the food until its excretion, allowing nutrient absorption ^3,4^. Peristalsis is a conserved mechanism in animals, from small planktonic crustaceans ^5^ to whales ^6^, and misfunctions in this process can lead to various pathologies in humans, such as gastroparesis, Hirschsprung disease, achalasia and others ^7,8^. Peristalsis can be regulated by various factors such as diet or microbiome composition ^9,10,11^, and is triggered by ICCs, cells with a pacemaker activity ^3,12,13^, expressing the Ca2+-activated Cl-channel ANO1/TMEM16A, which has been shown to be a selective marker for this cell type in mammals ^14^, while in *Drosophila* a marker for peristalsis has not been identified yet. To date, no other self-generated (i.e. pacemaker) contractions have been described in the gastrointestinal system other than peristalsis and segmentation (a peristalsis functional variant) ^4^, both driven by ICCs activity. Here, we uncover and characterize a novel mechanism by which the gut is able to produce strong muscular contractions independent from peristalsis. We show with direct imaging evidence that this activity, which we named “superstalsis”, is linked to visceral longitudinal muscle activation and is maintained for hours when the gut is excised from the rest of the body, reflecting a pacemaker function. We see that superstalsis is present *in vivo* in standard diet conditions and is significantly modulated in periods of altered intestinal homeostasis, such as bacterial gut infections or fasting. Moreover, we identify in *Subdued*, an ortholog gene of ANO1/TMEM16A, a marker for peristaltic activity in *Drosophila*, and show that its activation is specifically confined to circular visceral muscles. This work outlines a new scenario in the context of intestinal motility, where the interplay of peristalsis and superstalsis can be modulated by homeostatic and pathological cues, and it is achieved through the activation of specific subsets of visceral muscles.

## Characterization of superstaltic activity in isolated midguts

Observations on *ex vivo* fly intestines, led us to notice an unusually strong gut movement at a mildly acidic pH (**supplementary video 1**). We therefore systematically monitored extracted adult midguts free to move in saline solution at different pH values: we found that in a specific pH window, peaking at pH 5.5, the gut displayed strong contractions which could not be ascribed to peristalsis (**Fig. 1b; supplementary video 2 and extended data Fig. 1**), as these movements did not follow the peristalsis-typical distal contractions and relaxations along the longitudinal axis of the intestine. Instead, they would engage in strong angular “whip-like” movements and could travel up to 1.4 mm when allowed to freely move in the solution (**Fig. 1e; Fig. 1f and extended data Fig. 2**). Furthermore, these contractions followed a different pH activation curve (**Fig. 1d; extended data. Fig. 2**) and displayed an average angle change of 18.5 ± 5.7 ° (n=7 guts; **Fig. 1c; extended data Fig. 1**), unlike peristaltic events which are typically characterized by a null virtual angular change.

**Fig. 1.**
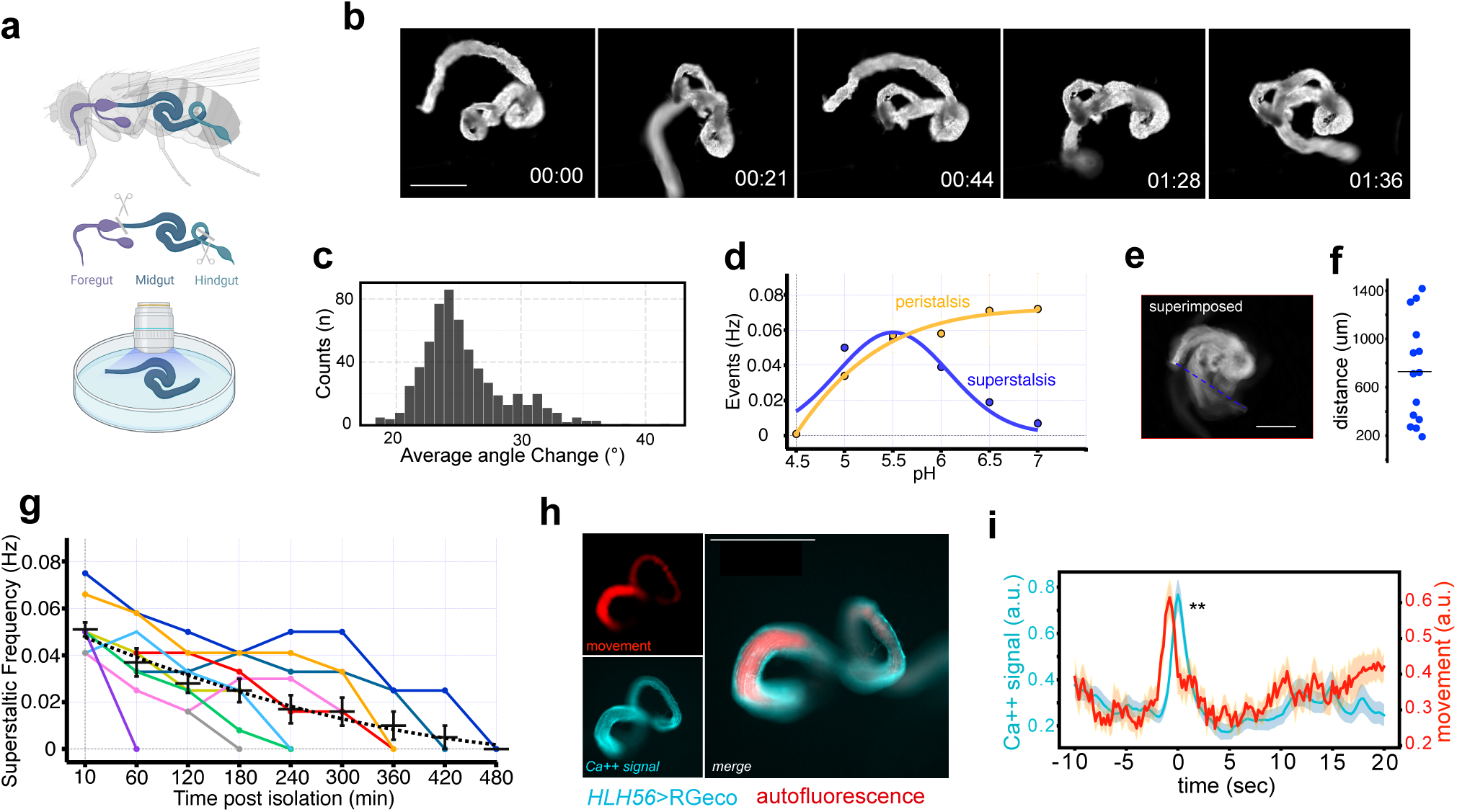
Superstalsis characterization. **a**, Cartoon depicting the experimental settings performed to record *ex-vivo* extracted midguts. **b,** Snapshots at different timepoints of supplementary video 2, showing the characteristic whip-like movement of superstalsis. Scale bar = 500 μm. **c,** Angular profile of recorded superstaltic in movie S2. **d,** Peristalsis and superstalsis pH-dependent profile of activation. Superstaltic events displayed a different pH activation window (pH 4.5, pH=5, pH=6.5, 0.019±0.007; pH 7, 0.007±0.003) compared to peristaltic ones (pH=4.5, 0.001±0.001; pH=5, 0.03±0.008 Hz; pH=5.5, 0.057±0.01 Hz; pH=6, 0.05±0.01 Hz; pH=6.5, 0.071±0.02 Hz; pH=7, 0.072±0.02 Hz). **e,** Image depicting superimposition of all the frames of the recorded video, through which we could calculate the maximum distance travelled by the gut at pH 5.5, summarized in **f**. Scale bar = 500 μm. **g**, Plot depicting superstalsis occurrence up to 7 hour post isolation (n=12 guts; each color corresponds to one gut). **h**, Representative snapshot from supplementary video 10, showing the two wavelengths (405 nm and 561 nm) used to compare movement and calcium signal during dual color imaging recordings. Scale bar = 500 μm. **i,** Dual color imaging traces showed a significant peak correlation between movement and calcium signals. Traces are mean (±SEM) of 7 movies (single Spearman correlation rho values and associated p-values are: r=0.75, p=0; r=0.41, p=0.0001; r=0.52, p=0; r=0.15, p=0.04; r=0.78, p=0; r=0.4, p=0; r=0.23, p=0.003)

**Fig. 2.**
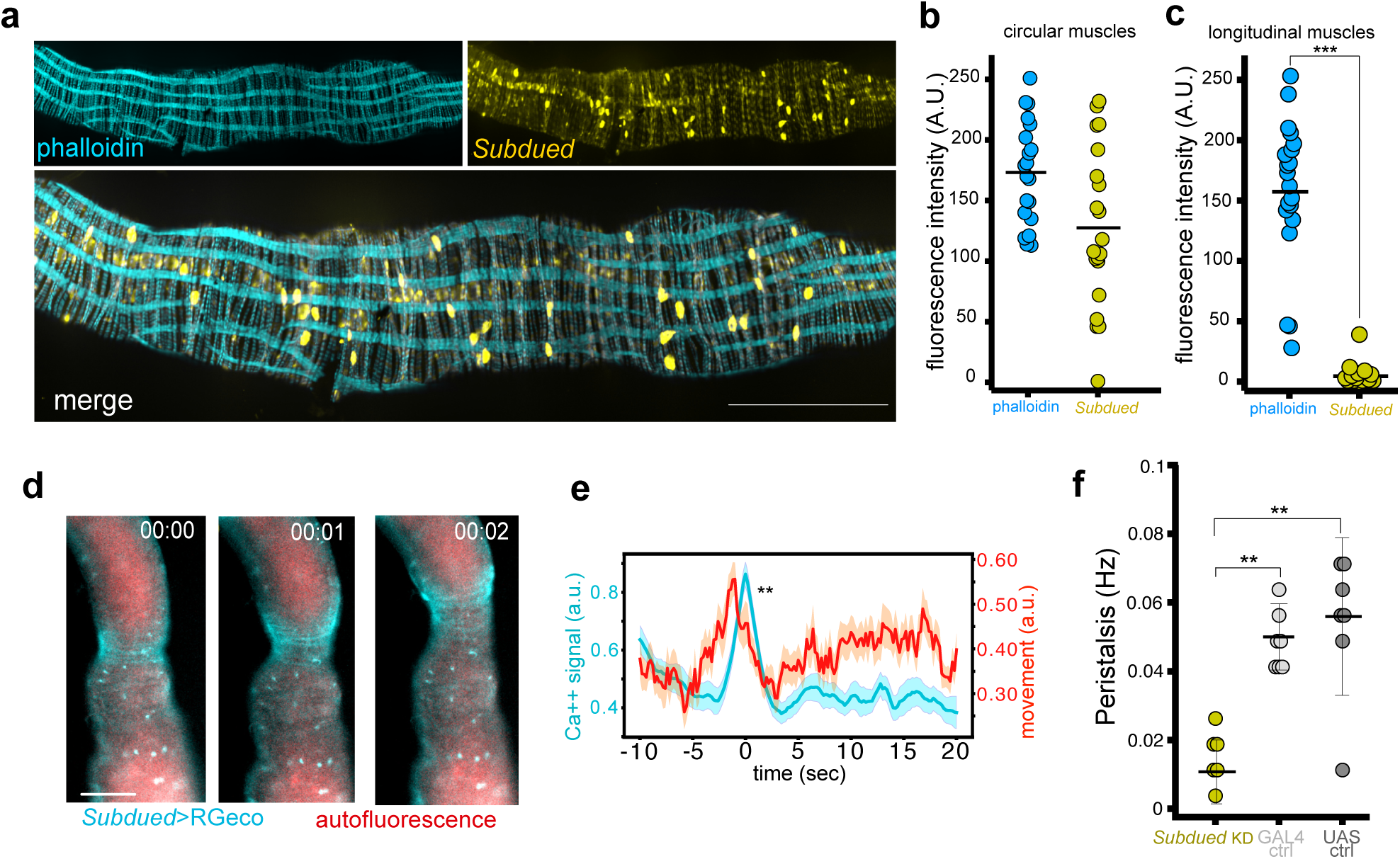
Subdued is a marker for peristalsis in *Drosophila*. **a**, representative confocal maximal intensity projection of stacked images of a posterior midgut expressing *Subdued*-Gal4>UAS-mCD8-GFP (yellow) and stained with phalloidin (light blue). In the merged channel colocalization is highlighted in grey (scale bar: 100 μm). **b** and **c,** Scatter dot plots of analyzed ROIs belonging either to circular (**b**) and longitudinal (**c**) muscles showing a subdued pattern expression confined to circular muscles. **d,** snapshots at different timepoints from supplementary video 8, showing the peristaltic wave coupled to an increase in fluorescence (scale bar: 100 μm). **e**, dual color imaging analysis of guts expressing *Subdued*-Gal4>UAS-RGECO confirmed a significant correlation between calcium increase and peristaltic events (n=9 guts; spearman rho and p values are respectively r=0.56, p=0; r=0.96, p=0; r=0.3, p=0.00015; r=0.45, p=0; r=0.56, p=0; r=0.45, p=0; r=0.47, p=0; r=0.57, p=0; r=0.23, p=0.0042). **f**, Knocking down subdued in muscle cells resulted in a significantly decreased peristalsis (subdued kd: 0.01±0.003 Hz, n=7 guts; Gal4 ctrl: 0.05±0.003 Hz; UAS ctrl: 0.055±0.008 Hz, n=7 guts) as summarized by this scatter dot plot (Genotypes are: *Subdued* KD = UAS-Subdued-RNAi>UAS-Dicer2/mef2-Gal4; Gal4 control = Mef2-Gal4>GD60000; UAS control = w1118/UAS-*Subdued*-RNAi;UAS-Dicer2)

At their activity peak (observed at pH=5.5), these movements were found to have a similar frequency to those measured during peristalsis at roughly 2/3 contractions per minute (**Fig. 1d; extended data Fig.2a**) and, although decaying with time, their occurrence was observed as long as 7 hours post isolation at room temperature without further oxygenation of the solution (**Fig.1g**), reflecting a pacemaker mechanism. Because of the aforementioned features and the relative high strength displayed by these contractions compared to peristaltic contractions, we termed this phenomenon “superstalsis”.

Superstalsis would occur several minutes after the gut’s exposure to the artificial saline solution, suggesting that the triggering cues would come from the novel solution permeating into the internal cavity of the gut rather than from the external environment (i.e. substances present in the circulating haemolymph; **extended data Fig. 2c**), in which case the activation would have been virtually immediate. Indeed, when flies were fed with the pH sensitive dye Bromophenol blue (BPB), the absence of superstalsis observed at time 0 correlated with a yellow colour in the copper cell region (already observed in ^15^; **supplementary video 3; extended data Fig. 2c**), while with the passing of time, the shift of this colour towards blue (and therefore to a pH> 3) was observed hand in hand with the appearance of superstalsis (**supplementary video 4; extended data Fig. 2C**). This confirms that superstalsis activation is triggered by changes in the gut lumen, where the food is processed.

To understand the visceral muscle contribution to the observed phenomena, we expressed the calcium reporter RGECO under the control of the pan muscular driver mef2 (mef2-Gal4>UAS-RGECO; **extended data Fig.3a**). Inspection of calcium imaging movies of isolated guts suggested that specific activation of the circular musculature was related to peristaltic events while an increased fluorescence in longitudinal muscles was associated to superstaltic events (**supplementary video 5; extended data Fig. 3b**). We therefore proceeded by addressing these two processes separately.

First, we expressed the calcium indicator RGECO selectively in visceral longitudinal muscles (HLH54F-Gal4>UAS-RGECO) and monitored the fluorescence intensity course during superstaltic events. To precisely determine the fluorescence increase signal in a three dimensional space, we performed dual color imaging, by exposing the gut to a 405 nm wavelength in order to simultaneously monitor the movement (see methods). As already evident by eye inspection, the analysis of the recorded traces showed a significant correlation between superstaltic contractions and longitudinal muscle-related calcium increase (**supplementary video 6; Fig.1h and i**), indicating a specific activation of this subset of muscles as actuators of the superstaltic activity.

To validate the relationship between longitudinal muscle activity and superstalsis, we investigated the possibility of selectively activating superstaltic movements by means of optogenetics. To mimic the observed physiological conditions, we expressed a recently engineered type of channelrhodopsin, CapChR2, which exhibits a selective permeability to calcium over other ions upon activation ^16^, in the longitudinal muscles.

Activation of CapChR2 induced superstaltic-like events in extracted midguts, while not generating peristaltic contractions (**extended data Fig. 4**). These observations show that superstalsis is selectively linked to longitudinal muscles in *Drosophila* midguts and can be driven independently of peristalsis.

## *Subdued* is a marker for peristalsis in *Drosophila*

In parallel, we aimed to unmask the molecular players responsible for peristaltic events. As, to date, a marker for peristalsis in *Drosophila* is not known, we reasoned that, since the peristalsis events observed in the *Drosophila* gut display similar anatomical and spatiotemporal features to those observed in other animals including mice and humans ^17, 18, 19, 20, 4^, this process might arise from a common ancestor gene of ANO1, the ICCs marker in mammals ^14^. In particular, one ortholog gene of the ANO1/TMEM16A family, *Subdued*, has been previously characterized in heterologous systems showing similar biophysical features to its mammalian counterpart ^21, 22^. We therefore examined the *Subdued* expression pattern in the *Drosophila* gastrointestinal tract. Immunostainings on extracted guts revealed an interstitial-like expression of *Subdued* positive cells in the midgut as well a strong expression in the hindgut and crop (**extended data Fig. 5a and 5b**). At closer inspection, we noticed that the *Subdued* signal was also found within the visceral muscles. Analysis of confocal stack images of midguts co-stained with phalloidin unmasked its expression specifically to the circular muscles, while virtually completely lacking from longitudinal ones (**Fig.2a, b and c; extended data Fig.5c, d and e**), confirming our preliminary observations on mef2>RGECO movies. Dual color imaging of RGECO-derived calcium signals showed a strong fluorescence increase, significantly coupled to the midgut peristaltic-related events when expressed under the control of *Subdued* (**Fig.2d and e; supplementary video 7 and 8**). Of note, *Subdued*-derived calcium signals were selectively associated with peristaltic, but not superstaltic events (**supplementary video 9**), indicating the existence of an independent regulation of these two activities. A strong *Subdued* activation concurrently with peristaltic contractions was also observed in the isolated crop (**supplementary video 10: extended data Fig. 6a, b and c**) as well as in the isolated hindgut (**supplementary video 11; extended data Fig. 6d**), where phalloidin-positive longitudinal muscles are absent ^23^, confirming the strong expression observed in confocal images (**extended data Fig. 5a**) and further validating the link between peristalsis, circular visceral muscles and pacemaker activity. Interestingly, as also seen in the hindgut, we noticed in some instances the appearance of calcium waves not leading to peristaltic contractions spreading throughout the midgut (**supplementary video 12**), suggesting a complex regulatory system which will need further investigations in the future. Finally, we evaluated the effects of knocking down *Subdued* on midguts motility profiles. The expression of *Subdued* RNAi confined to muscle cells resulted in a significantly impaired peristaltic activity compared to controls (**Fig. 2f**), confirming its functional role. Altogether these observations indicate that *Subdued* is a marker for peristalsis in *Drosophila*.

## *In vivo* monitoring of superstalsis in standard and altered gut conditions

We next explored the presence of superstalsis *in vivo* and in the context of an altered intestinal homeostasis. As previous studies showed a correlation with gastrointestinal muscle activation and bacterial infections^24^, we also tested a potential involvement of superstalsis in the context of gut infection, as well as in standard physiological conditions. We opted for *Pectobacterium carotovorum (Ecc15)*, a bacterial strain widely used in the context of *Drosophila* gut infection, known to activate the host immune system ^25^, as well as to induce increased defecation ^26^.

Because monitoring of the gut is shaded by the cuticle in adult flies, we focused on a preparation in which only the middle midgut was exposed to the solution, while still being connected to the rest of the body (**Fig. 3a**). For these experiments, midguts were exposed to *Drosophila* Schneider medium (pH ≃ 6.5), which is commonly used to record gut activity *in vitro* or to prepare cultures of the fly intestines ^17^. To test a context of non-pathological gut altered homeostasis, we also monitored guts of flies starved for 24 hours (**Fig. 3b**), as starvation has also been shown to alter the gut physiological conditions ^27,28,29^. Interestingly, we found a significant increase of superstalsis both upon *Ecc15* infection (**supplementary video 13**) and after 24 hour starvation (**supplementary video 14**) compared to controls (**supplementary video 15; Fig. 3b and c**). In contrast, the peristaltic activity was slightly increased upon bacterial infection and, to our surprise, almost absent in starved flies (**Fig. 3d**), outlining a complex gut motility regulation driven by diverse alterations in the gut homeostasis.

**Fig. 3.**
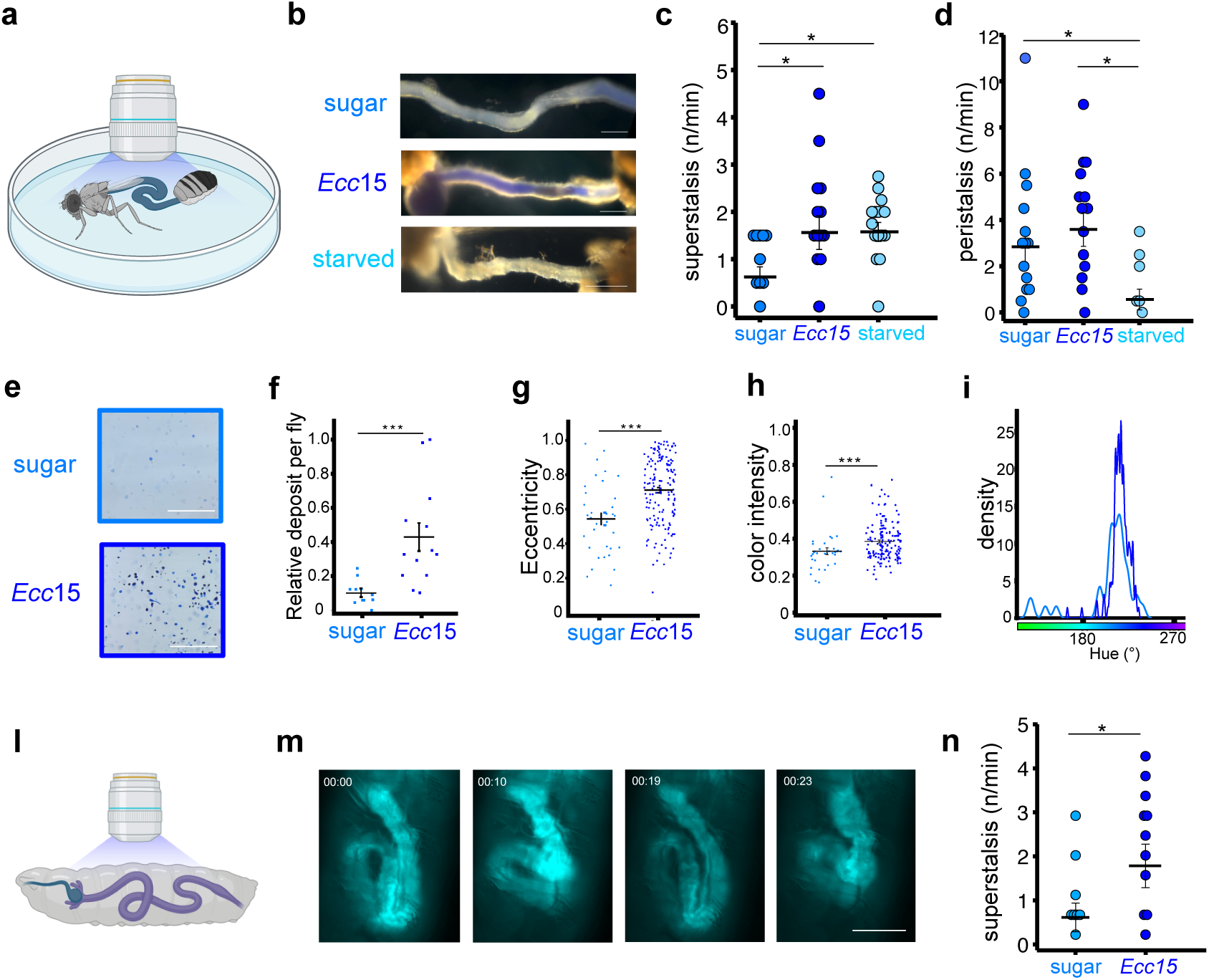
Superstalsis modulation and monitoring *in vivo.* **a**, cartoon showing the experimental settings used to monitor exposed guts in adults. **b**, representative images of monitored guts upon 2 hours fed with sugar and BPB, Ecc15 and BPB or fasted for 24 hours. A shift is noticeable in the BPB in infected flies compared to controls indicating an altered pH (scale bar: 400 μm). **c,** superstaltic events increased significantly upon both *Ecc*15 infection (0.03±0.005 Hz, n=16 guts) and 24 hours fasting (0.03±0.003 Hz, n=16 guts) compared to controls (0.01±0.002 Hz, n=16 guts). **d,** peristaltic events on the other hand did not significantly change in infected flies (0.06± 0.01 Hz), while were drastically reduced upon 24 hours fasting (0.01± 0.004 Hz) compared to controls (0.05± 0.01 Hz). **e,** representative images of adult flies fecal output after 2 hour exposure to sugar and *Ecc*15 (Scale bars: 10 mm). *Ecc*15 caused an increased number of deposits (**f**) as well as significant changes in the shape (**g**) andintensity (**h**) of the excreta, associated with a shift in their color (**i**). **l**, cartoon depicting schematized experimental setting for imaging whole guts in intact larvae. **m** snapshots of different timepoints from supplementary video 16, documenting the existence of superstalsis in vivo. Scale bar: 500 μm. As shown in the scatter plot in (**n**) superstaltic events were present in standard diet conditions (0.01±0.004 Hz, n=13 larvae) and were significantly enhanced upon 24 hour infection with *Ecc*15 (0.03±0.007 Hz, n=14 larvae)

In parallel, we performed the fecal output assay, a reliable behavioral assay to monitor changes, among which the pH, associated with altered gut conditions. Through this assay we could gather various information about intestinal homeostasis, such as intestinal transit and acid-base balance in adult flies^30^ (**Fig. 3e**). Analysis of deposits of adult flies fed for 2 hours solution with the pH sensitive dye BPB added to either sugar or *Ecc*15 revealed differences in several parameters examined. In fact, besides the increased number of deposits (as already shown in ^26^; **Fig. 3f**; sugar=0.1±0.02; *Ecc*15=0.42±0.08), we found differences also in the shape (**Fig. 3g**; sugar=0.54±0.03; *Ecc*15=0.71±0.01) and intensity (**Fig.3h**; sugar=0.33 ± 0.019; *Ecc*15=0.38±0.008) of the deposits of infected flies, as well as a shift in their colour (**Fig. 3i**). Importantly, these results show a strong relation between gut motility (increased number and total amount of deposits) and pH shift in the gut lumen (fecal output density and colour), validating our observations on isolated and exposed guts.

Lastly, in order to monitor superstalsis in a full physiological context, we switched to fly larvae, as, since they are visually accessible, we could evaluate the gut motility in intact animals (**Fig. 3l**). To better visualize gut movements, we added the fluorophore Alexa488 to the sucrose or *Ecc*15 solution. Strikingly, superstalsis was observed in standard diet conditions (**supplementary video 16; Fig. 3m**) and, as similarly observed in adults, *Ecc*15 gut infection significantly increased superstaltic events (**Fig.3n**), indicating the existence of superstalsis in basal conditions *in vivo*, and confirming our previous results.

## Discussion

The gastrointestinal system relies on complex molecular pathways aimed to ensure a correct metabolic function and, at the same time, act as gatekeepers in order to prevent harmful events driven by exogenous substances ^31,32^. As such, the intestine has to face, interpret, and respond to many factors, including food molecules, chemicals and microorganisms which come in contact with it. In this context, as for other involuntary body mechanisms such as the heartbeat or breathing, pacemaker activities guarantee a correct function of the system, overruling voluntary actions. To date the only reported enteric pacemaker activity is the one driven by the ICC activity, leading to peristalsis, segmentation or the migrating motor complex ^33,4^. This study describes and characterizes superstalsis as a previously unreported self-generated slow-wave activity in the fly intestine. We here show how this novel pH-dependent pacemaker slow-wave activity is enhanced upon conditions of altered gut homeostasis (i.e. bacterial infection) *in vivo*. Our findings unveil the existence of a connection between superstalsis activation and events (either pathological or homeostatic) leading to an altered gut pH profile, such as intestinal-related diseases ^34^ or microbiome composition ^35,15^. Indeed, if on the one hand the gut microbiome has been shown to affect the fly gut pH profile both in adults^15^ and in larvae^36^, on the other hand the intestinal bacterial composition is itself subject to dynamic changes, driven by many factors, such as bacterial infections^15,37^, changes in the environmental niche^38,39^, diet composition ^40,41^, age ^42^ and starvation^29,43^. What emerges is a scenario in which a variety of cues derived from an altered gut environment lead to the specific activation of circular (modulating peristalsis) or/and longitudinal muscles, activating superstaltic contractions.

Our evidence shows the enteric ability to self-regulate its own motility profile through the specific balance of these functionally parallel and anatomically orthogonal mechanisms. The observation that in basal condition this equilibrium is shifted towards peristalsis, while upon altered homeostatic events it is transposed towards superstalsis, indicates peristalsis to be responsible for physiological food processing and nutrient absorption, while superstalsis rather being reinforced to potentially prevent, through an efficient emptying of the gut lumen, harmful conditions such as increased acidity or bacterial infection. This would explain the high strength related to superstaltic contractions compared to peristaltic ones, and its particular pH activation window.

The involvement of visceral longitudinal muscles has been previously reported in mammals in relation to emesis^44^. Several considerations make the involvement of superstalsis in vomiting behavior unlikely: first, the Retrograde Giant Contractions (RGC) associated with emetic episodes in mammals have been shown to be under extrinsic control ^45^, therefore not reflecting a pacemaker activity. Second, a recent study identified the P4 pump of the foregut as the main effector of vomiting in *Drosophila* ^46^, a part of the gastrointestinal system not included in our *ex vivo* preparations nor monitored during recordings on exposed guts.

Molecular mechanisms leading to longitudinal muscle activation through the activation of the Dual Oxidase (DUOX) pathway upon bacterial gut infection was previously reported in *Drosophila*^24,26^. Although we also documented an increased superstalsis upon bacterial infection, we have also seen its increase upon fasting, suggesting the presence of parallel pathways leading to a similar activation of this muscle subset. Furthermore, because superstalsis was observed in standard physiological conditions *in vivo*, we speculate that its activation might arise from different cues besides chemical, such as thermal or mechanical: especially taking into account the anatomical disposition of longitudinal and circular visceral muscles being perpendicular to each other. It would appear to be a more efficient scenario if each muscle subset could give their own angle contribution to the overall gut motility profile, allowing more power and control for the gut to move itself. This feature might be fundamental in an anatomical context where an intricate tubular system, with various turns and U-loops, is confined to a limited space.

While studying the different sources of peristaltic and superstaltic gut movements, this study identified *Subdued* as a marker for peristalsis in *Drosophila*. Previous work identified *Subdued* as a TMEM16A ortholog gene and characterized it in a heterologous system, unveiling similar biophysical properties to the mammalian TMEM16 channel family^21, 22, 47^. *Subdued* was found to be expressed in the intestine and knockout flies for this gene displayed an increased mortality upon bacterial gut infections ^21^. These findings support our evidence, as we reported that *Subdued* is expressed specifically in circular visceral muscles and its activation is coupled to peristaltic events, as well as *Subdued* RNAi confined to muscle cells is decreasing peristaltic activity.

Overall, our findings unveil a scenario in which two types of contraction with pacemaker activity are present and can be independently modulated by the gut itself as a consequence of specific pathophysiological intestinal changes. Future studies will have to clarify the molecular pathways leading to superstalsis and a potential presence of superstalsis in mammals, but taking into account the existence of a clear anatomical separation (unlike in flies) between circular and longitudinal muscles by the Auerbach (i.e. myenteric) plexus ^48^, it is tempting to speculate that throughout evolution these mechanisms have been further refined independently in order to accomplish their specific distinct functions, allowing the gut to finely regulate its responses.

## Materials and methods

### Animal husbandry and genetics

Flies were reared on standard cornmeal food at 25°C (60% humidity) under a 12h dark/light regime. 3 to 7 days old flies were used, unless otherwise indicated. The following stocks were used: w1118 (BDSC:3605); UAS-GCaMP6m (BDSC:42748); UAS-jRGECO1a (BDSC: 63794); Mef2-Gal4 (BDSC: 27390); Subdued-Gal4 (BDSC: 97156); HLH54F-Gal4 (BDSC: 91633); UAS-mCD8-GFP (BDSC: 32185); UAS-Dicer2 (BDSC: 24651); UAS-Subdued RNAi (v37472); GD controls (v60000); UAS-CapChR2 (BDSC:98422). Crossing genotypes are indicated throughout the manuscript.

### *Ex-vivo* imaging and analysis of gastrointestinal organ activity

To monitor the ex-vivo enteric pacemaker activity, adult male flies were starved for 2 hours, anesthetized on ice and subsequently gastrointestinal organs (either midgut, hindgut or crop) were extracted in saline containing (in mM): 130 NaCl, 5KCl, 2MgCl2, 2CaCl2, 5 HEPES (≈ 270 mOsmol/kg). The pH was adjusted to the desired value by adding NaOH or HCl and measured with a pH-meter Knick Calimatic 765. Midguts were isolated from the rest of the gastrointestinal parts, transferred in a petri dish containing the same solution and imaged free to move. Where indicated, Drosophila Schneider medium (Capricorn Scientific; SCH-1000ML) was used as a dissection and recording solution. To monitor the gut activity, we used a Nikon Widefield Bright Field equipped with a sCMOS PCO edge camera (2048×2048 pixels, 6.5 µM pixel size), either in phase contrast modality (to monitor gut movements; 5 Hz sampling rate, 2 min length), or in EPI Fluorescence modality (for Calcium and Voltage imaging experiments; 5 Hz sampling rate). Either a Nikon 4x Plan Apo Air (NA=0.2) or a Nikon 10x Plan Fluo Air (NA=0.3) objectives were used. To quantify superstalsis frequency, recorded movies (2 min long) were imported and analyzed in NOSA^49^. Angular gut movements on the XY axes were classified as superstaltic events each time the gut displayed an angular movement ≥ 10° on the XY axes, or when the gut translational displacement on the XY axes exceeded at least half of the diameter of the gut itself. An n=1 superstaltic event comprehended both the contraction and relaxation events of the muscular movement. Peristaltic events were quantified manually and the counted numbers confirmed with NOSA.

### Optogenetics

Midguts were isolated from 3-7 days old flies expressing the Calcium permeable channerhodipsin CapChR2^16^ in visceral longitudinal muscles (HLH54F-Gal4>UAS-CapChR2), transferred to a dish containing Drosophila Schneider Medium and monitored with a Nikon Widefield Bright Field equipped with a sCMOS PCO edge camera with a Nikon 10x Plan Fluor Air objective. To achieve channelrhodopsin activation, guts were exposed to a 488 nm LED light. As negative control, the same guts were exposed to a 647 nm LED light.

### Confocal imaging and analysis

Extracted guts were fixed for 20’ at Room Temperature (RT) in 0.3% Paraformaldehyde (PFA), washed three times in 0.3% PBST and incubated with the antibodies at 4°C overnight. After 3 washes in PBST, samples were then mounted and imaged the next day. Stacked Images were captured using a Spinning Disk confocal CSU-X Microscope (Nikon), equipped with a EMCCD camera (iXon3 DU-888 Ultra, 1024×1024 pixels) Nikon Plan Fluor 40x oil immersion objective (NA=1.3). Mounted Samples were excited using a 488, 561 nm and a 647 nm lasers and Fluorescence emission was detected using GFP, mCherry and Cy5 filters and images were stitched together to capture the whole CNS structures of interest (z-stack = 0.7 µM). Phallodin stainings were performed by incubating fixed samples at 4°C with either FITC-coupled Phalloidin (SIGMA; P5282) or phalloidin-iFluor(TM) 647 conjugate (AAT Bioquest; ABD-23127) at 1:500 for 45’. To quantify *Subdued* expression in visceral muscles, maximal intensity projections of stacked images of midguts co-stained with Subdued-mCD8-GFP and Phalloidin were analysed. First, ROIs corresponding to longitudinal or circular muscles in the phalloidin channel were selected. Afterwards, the same ROIs were applied to the same image on the Subdued-mCD8-GFP channel and the fluorescence intensity was quantified.

### Intestinal Movement Tracking and correlation of motion with Calcium Dynamics

Time-lapse movies of fly intestines were processed using IntestiTrack, a custom Python script implemented in Jupyter Notebook (https://github.com/ngimber/Intestinal-Dynamics-Toolbox). Raw images were segmented by Otsu’s thresholding (49), followed by dilation, and skeletonization. The resulting skeletons were pruned to remove spurious branches and intestinal movement was tracked across timepoints. To visualize the dynamics, skeleton overlays of consecutive frames were generated. For quantitative analysis, the skeleton coordinates were used to compute angular changes over time. Histograms of mean angle change per time step were plotted to quantify the distribution of movement dynamics.

For dual color imaging, intestines expressing UAS-RGECO in the desired cell type were monitored at 5 Hz by exposing them to alternating LED pulses at 561nm and 505 nm (the latter to detect the motion through autofluorescence). Dual color image data were analyzed using ‘CaMotion’, a custom Python script implemented in Jupyter Notebook (https://github.com/ngimber/Intestinal-Dynamics-Toolbox). To quantify temporal movement, intestines were segmented using Otsu’s thresholding (49). Then the Pearson correlation coefficient was computed between each frame and the corresponding frame acquired one second earlier, providing a measure of motion dynamics (movement = 1 – correlation coefficient). Peaks of calcium transients were identified by detecting local maxima in fluorescence intensity traces using the ‘find_peaks’ function of SciPy^51^. Traces of both channels around detected calcium peaks were temporally aligned such that the peak maximum was defined as t = 0, enabling averaging across events. Traces of movement and calcium transients were normalized to a 0–1 range before plotting.

### Oral infection

Oral infection of adult flies was performed as adapted from ^52^. Briefly, flies were kept in empty vials at 29°C for 2h prior to infection. *Pectobacterium carotovorum* (*Ecc15*) was grown in LB medium overnight at 30°C. The cultures were then centrifuged for 15 minutes at 3500 g and 4°C and diluted in PBS to an OD600 of 400. This bacterial suspension was mixed 1:1 with a 5% sucrose and 1% bromophenol blue solution, and 150 µL of the mixture pipetted onto vials containing standard food completely covered by filter paper. For oral infection of larvae, filter paper was soaked in 150 µL of the same mixture used for adults supplemented with Alexa-488 fluorophore (1:100, ThermoFisher A13201).

### Exposed gut activity recording and in vivo monitoring of superstalsis in *larvae*

To monitor superstalsis in adults, wild-type (w1118) flies of both sexes, 3-7 days old, were collected, starved for 2 hours and subsequently exposed to a 2 hours oral infection as described above. For the starvation group, flies were starved for 24 hours in total before recordings.

Flies were then briefly anesthetized on ice and subsequently submerged in a petri dish containing 2 mL of Drosophila Schneider Medium (Capricorn Scientific; SCH-1000ML) at RT. Once legs and wings were removed, a cut was performed between the thorax and the abdomen to expose the midgut to the solution. The upper and lower part of the body were then fixed to the base of the petri dish (previously coated with agarose) with small needles. The gut was then imaged on a Nikon Widefield Bright Field equipped with a Nikon DS-Ri2 color CMOS camera (712 x 712 pixels) and a 4x Plan Apo λ Air (NA 0.2 WD 20,000 µm) and imaged at 5 Hz for 2 min.

To monitor gut movements in larvae, wild type (w1118) 3rd instar larvae were collected, starved for 2 hours and exposed to the filter paper containing sugar or *Ecc15*, as described above. Larvae were then immobilized on a 4% Low-Melting Agarose (Euromedex LM3; ref:1670) solution onto a coverslip and imaged with 488 nm LED exposure for 2 min with a Nikon Widefield Bright Field equipped with a sCMOS PCO edge camera using a Nikon 4x Plan Apo Air (NA=0.2) objective.

### Defecation assay

Defecation analysis of adult wild-type flies (2–6 days old) was performed as described before^30^. In brief, flies were subjected to oral infection with Ecc15 or a sugar control for 2 hours at 30 °C. Following exposure, vials were imaged digitally for further analysis of fecal output using a custom-written Python-based image processing pipeline implemented in Jupyter Notebook.

For each image, a region of interest (ROI) with a 700 px radius was randomly selected. Fecal deposits were detected by clustering pixel values using a Gaussian mixture model. False positives were manually removed. For each condition, the number of fecal deposits per fly was quantified and normalized to the minimum and maximum values within the dataset. Deposit size was defined as the total number of pixels associated with each fecal spot. Shape was quantified using eccentricity, where a value of 0 corresponds to a perfect circle and 1 to a straight line. For intensity and color characterization, mean red, green, and blue channel values (RGB) were converted to hue, saturation, and lightness (HSL). Lightness was used as a proxy for fecal density, and hue values were used for visualization of color variation between experimental groups.

### Statistics

Data are presented as standard error of the mean (SEM) unless otherwise indicated. To check statistical significance among different conditions tested we used the unpaired t-test for parametric data and two-sided Mann-Whitney for non parametric data. P-values <0.05 were considered statistically significantly different and annotated as: p<0.05 =*; p<0.01=**; p<0.001=***.

## Acknowledgments

The authors thank Irene Miguel-Aliaga for comments and advice on preliminary data; Bruno Lemaitre lab for *Ecc15*-GFP bacteria (used for preliminary experiments in larvae); members of the Owald lab for help throughout the project; the Advanced Medical BIOimaging Core Facility of the Charité-Universitätsmedizin Berlin (AMBIO) for support in acquisition and analysis of the imaging data.

## Funding

European Research Council (ERC) grant ‘SimpleMinds’ (101088502) to DO. Deutsche Forschungsgemeinschaf (German Research Foundation) DFG grant (52455519) to NPP.

Studienstiftung des deutschen Volkes e.V. doctoral scholarship to ISB.

## Author contributions

Conceptualization: NPP, DO

Methodology: NPP, ISB, PD, NG, ER, AE

Investigation: NPP, ISB, PD, NG, ER, AE

Visualization: NPP, ISB, PD, NG, ER, AE

Funding acquisition: NPP, DO

Project administration: DO, II, JS

Supervision: NPP, DO, II, JS

Writing – original draft: NPP, DO

Writing – review & editing: NPP, DO, II, ISB, PD, NG, ER, AE, JS.

## Extended data

**Fig. S1.**
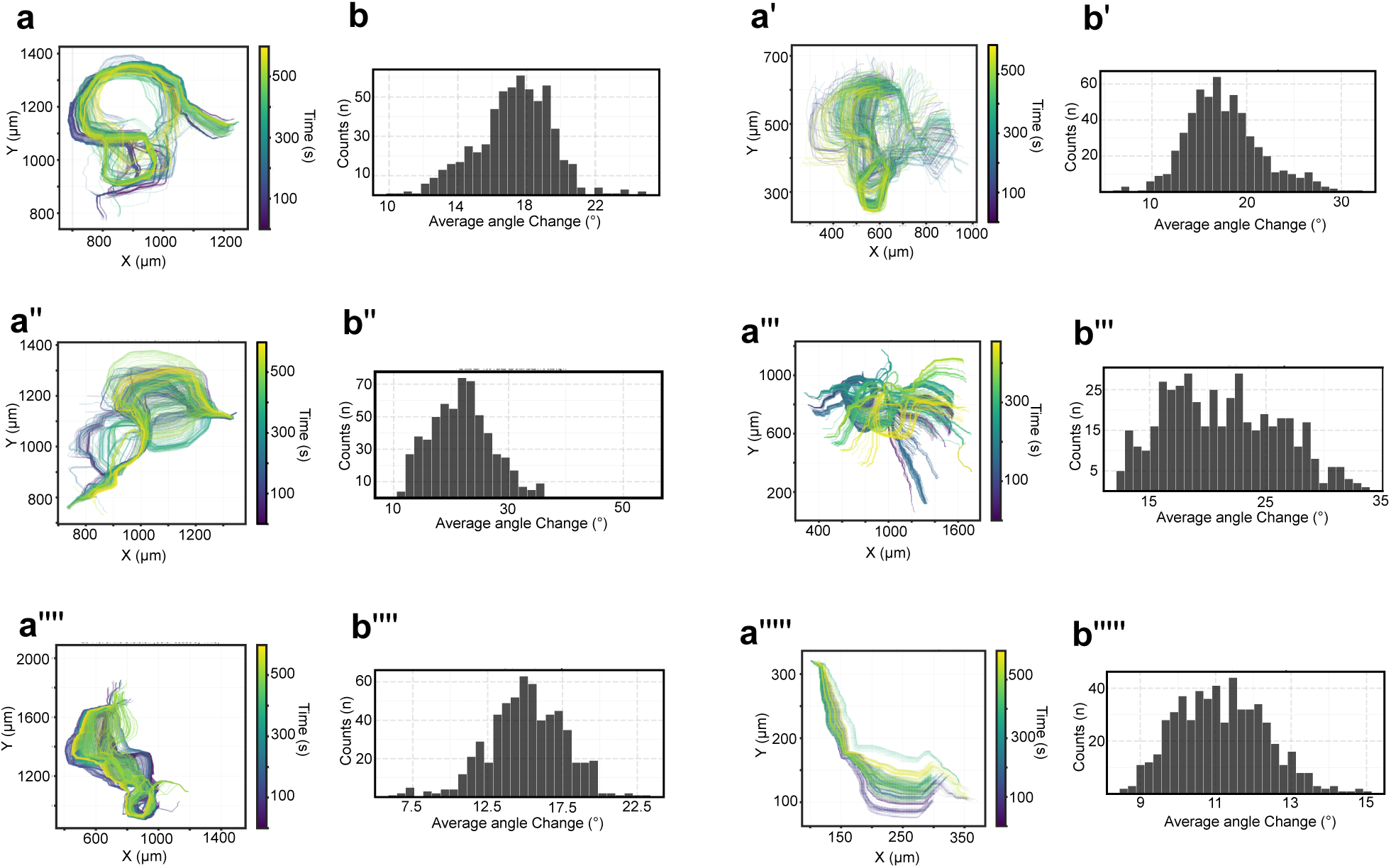
Superstalsis spatiotemporal characterization. Color coded time motion profiles (**a**) and Angular profiles (**b**) from 6 representative extracted midguts recorded for 2 min.

**Fig. S2.**
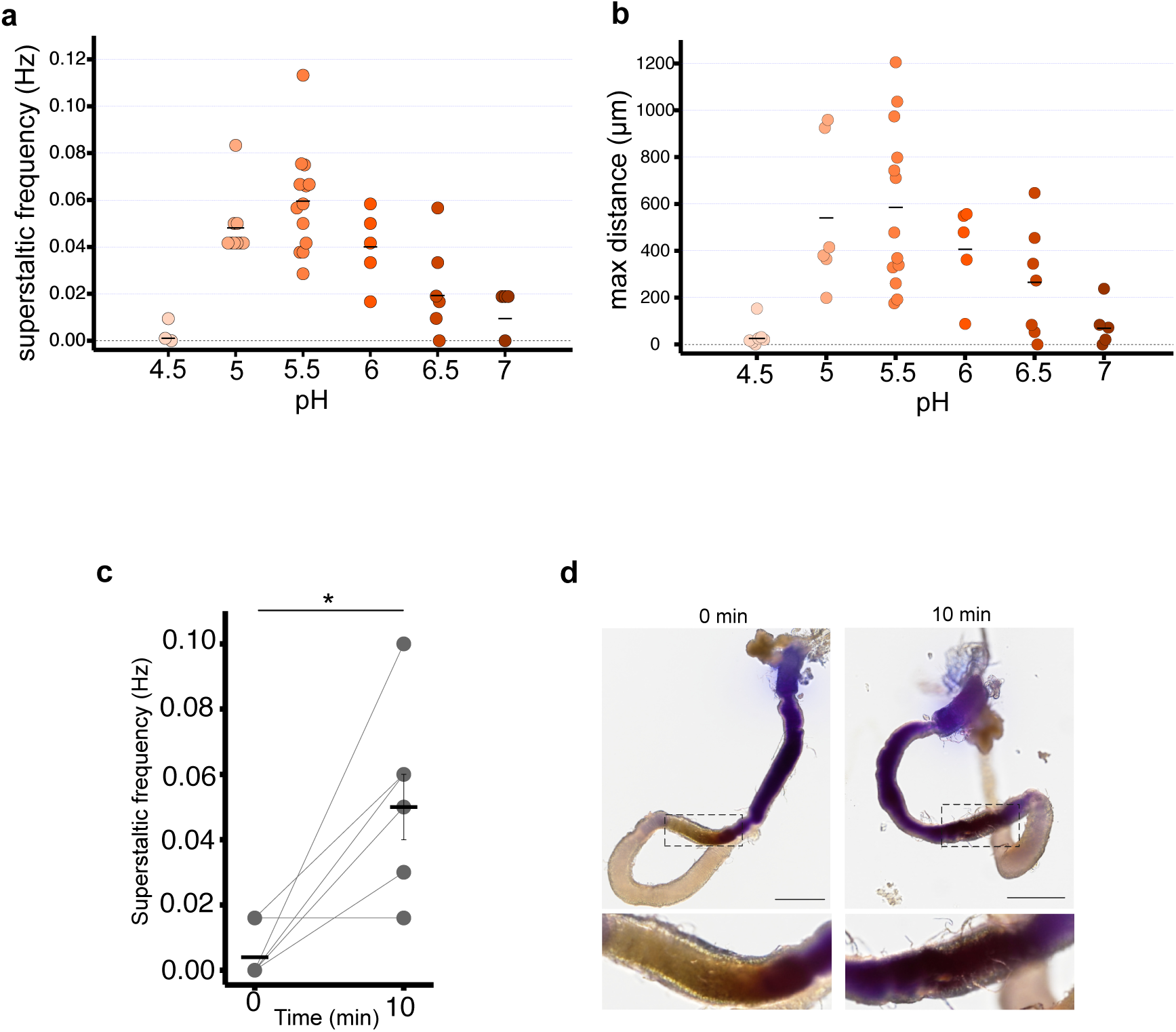
Superstalsis dependence on pH. **a**, Scatter plot showing the pH window in which superstalsis is active, peaking at pH 5.5. Each dot represents a different gut recorded for 2 min. On the same recorded movies we calculated the maximum distance covered by the superstaltic events during the time of the recording and, as shown in the plot in (**b**) there was a correlation between frequency and strength (distance) of the contractions. **C,** plot quantifying superstaltic frequency on guts recorded at time 0 and after 10 minutes (same gut) of exposure to the artificial saline solution, indicating the exchange of the novel solution into the gut lumen. **d,** fly gut isolated from a BPB fed fly showed a change in color corresponding to the copper cell region (highlighted) at time 0, associated to null superstalsis, and after 10 min, associated to peristalsis occurrence (scale bars: 500 μm).

**Fig. S3.**
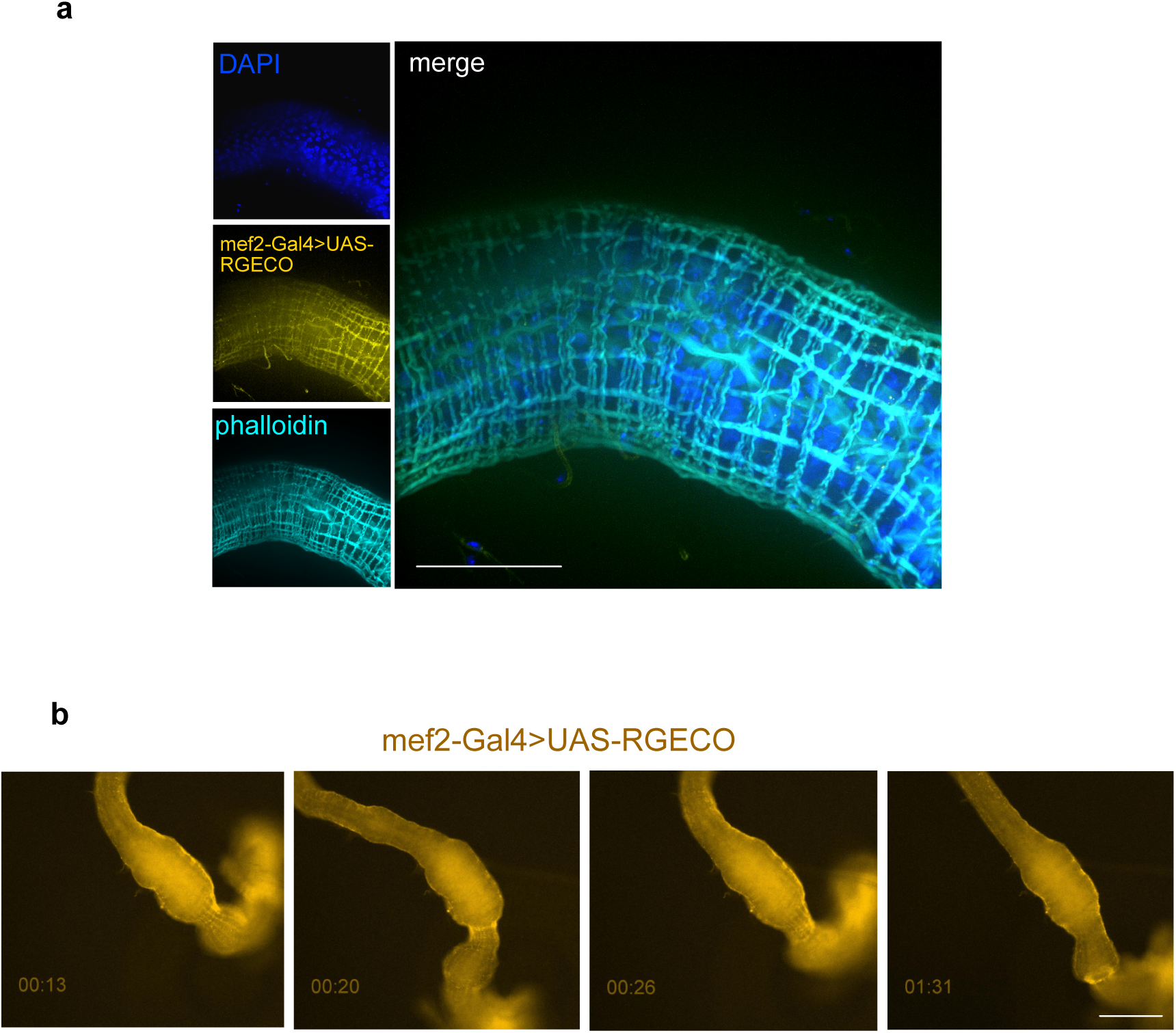
Pan-muscular Calcium imaging. **a**, confocal image of a midgut expressing the calcium reporter RGECO under the control of the pan-muscular driver mef2 fixed and subsequently stained with phalloidin shows a complete colocalization of mef2 and phalloidin signal (scale bar: 50 μm). **b**, snapshots from supplementary video 5 at different timepoints. Is noticeable the increased fluorescence associated to circular muscles during peristaltic events and to longitudinal ones during superstalsis (scale bar: 200 μm).

**Fig. S4.**
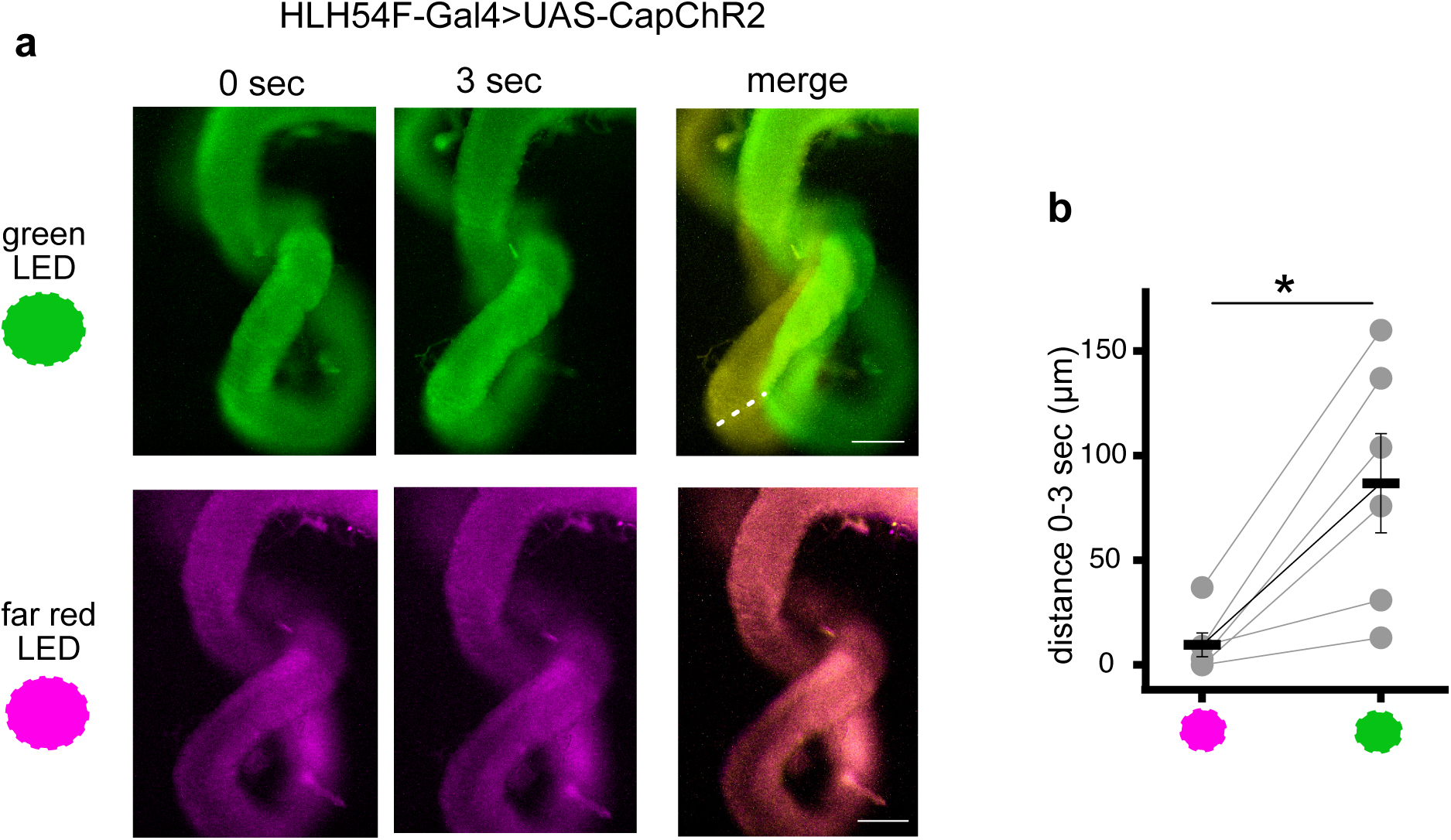
Optogenetic activation of superstalsis. **a**, Snapshots of adult fly guts expressing the calcium permeable CapChR2 channelrhodopsin in longitudinal muscles (HLH54F-Gal4>UAS-CapChR2) at time 0 and after 3 second of exposure to either 488 nm LED (upper panel; green) or 647 nm LED (bottom panel; magenta). It is evident a superstaltic-like motion after 3 seconds of 488 nm LED exposure (activating CapChR2), while no movements were observed in the same gut upon 647 nm LED exposure. **b**, plot representing the position difference of the same gut measured at time=0 or 3 seconds after exposure of either 647 nm LED (left) and 488 nm LED (right). Activation of longitudinal muscle through calcium permeation (green light) induced a significant contraction (86.8±23.7 μm) compared to control illumination (far red light; 9.67±5.7 μm). Scale bar: 100 μm

**Fig. S5.**
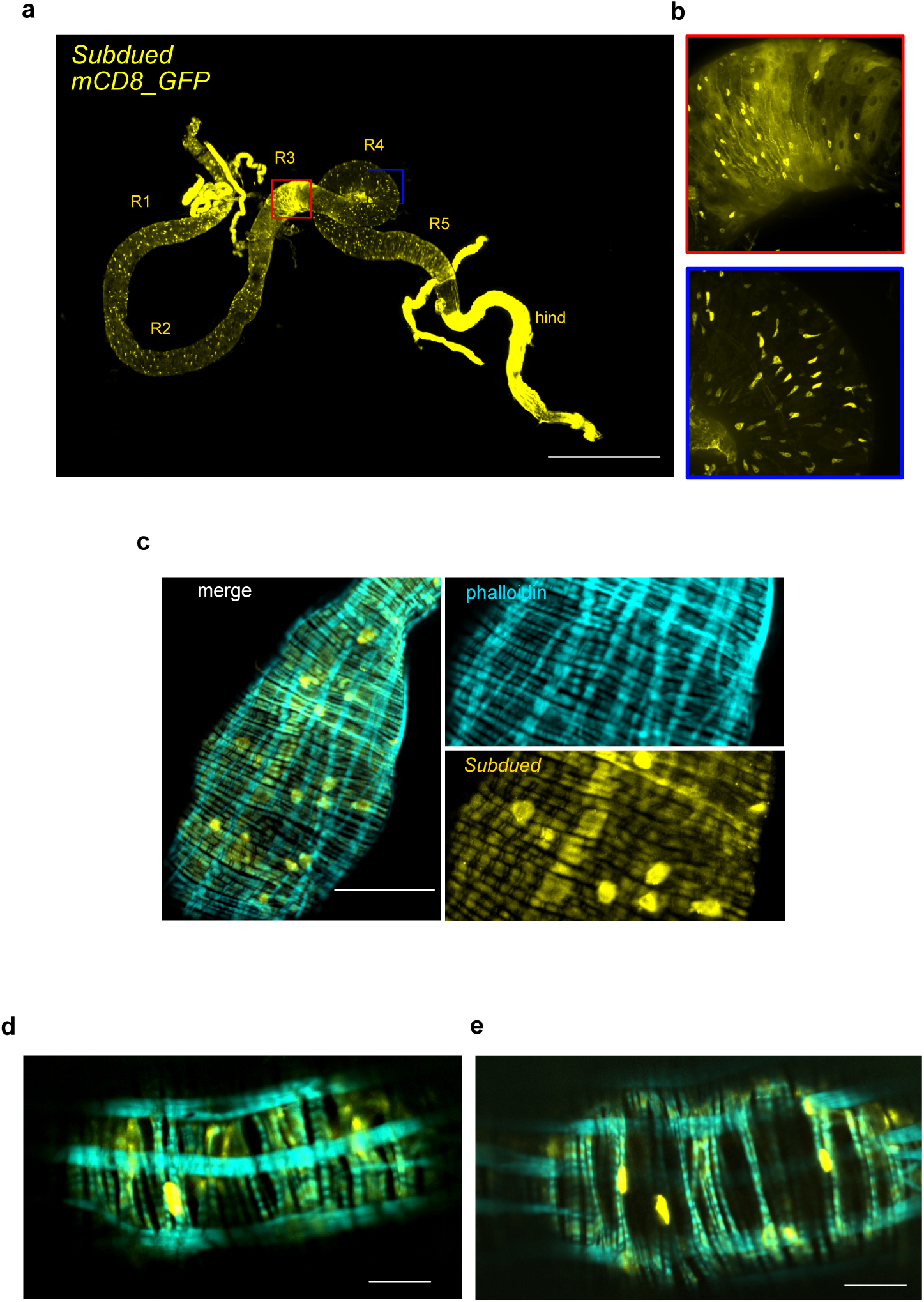
*Subdued* expression profile in adult intestines. **a**, Confocal image representing the expression profile of *Subdued* throughout the whole adult drosophila intestine (*Subdued*-Gal4>UAS-mC8-GFP). Midgut subregions are labelled from R1 to R5 (as described in Buchon et al, 2013). The R3 and R4 regions are highlighted in the insets in (**b**). Scale bar: 500 μm. **c**, example of a midgut expressing *Subdued*-Gal4>UAS-mC8-GFP and stained with phalloidin highlights the absence of gfp signal from longitudinal muscles. Scale bar: 50 μm. (**d**) and (**e**) Zoomed subportions from the gut image shown in fig. 2 evidence as well how the presence of the *Subdued* signal is confined to circular muscles. Scale bars: 20 μm.

**Fig. S6.**
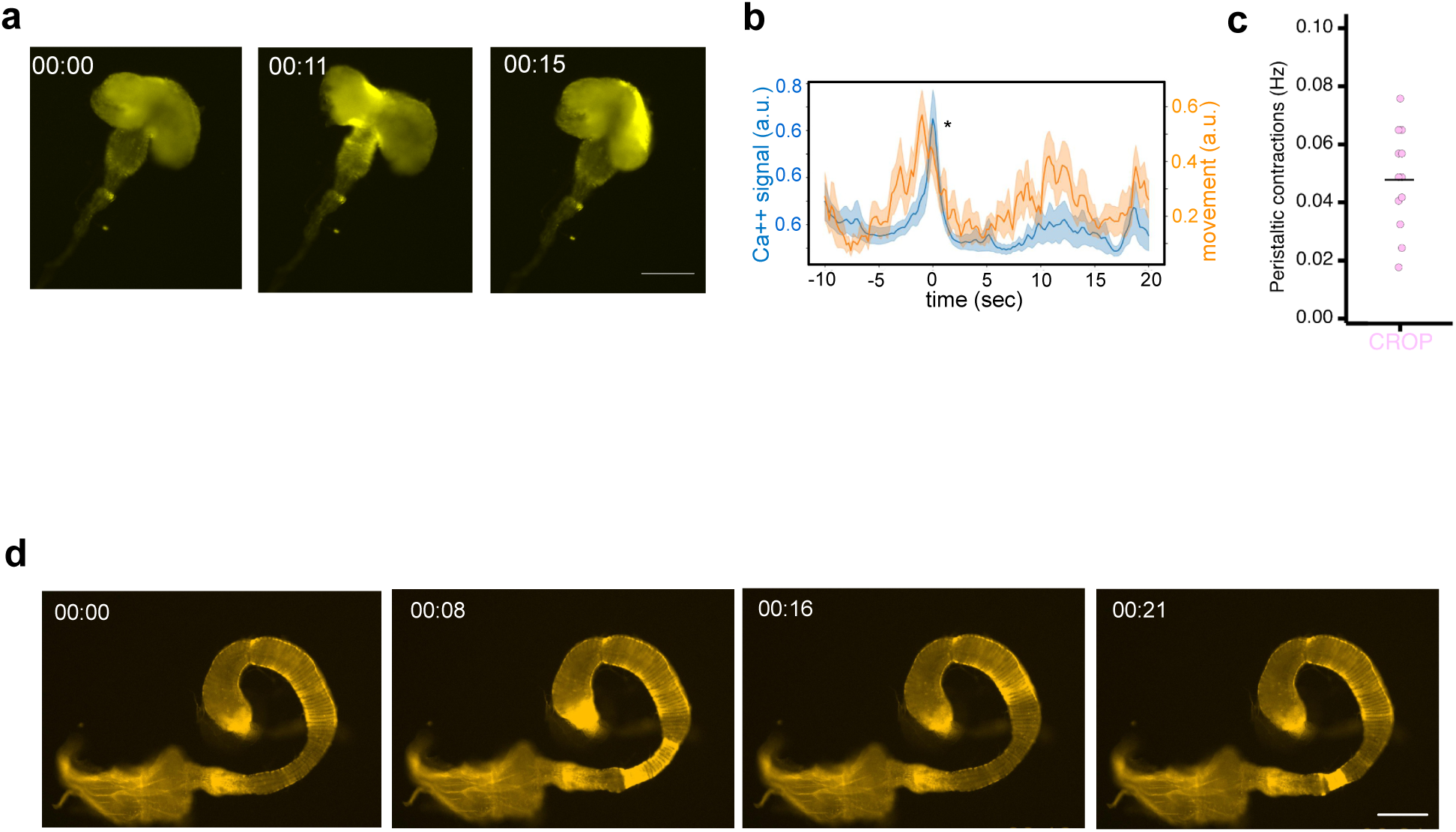
Subdued expression in the crop and the hindgut correlates with peristalsis. **a**, snapshots of supplementary video 11 at different timepoints showing the increase in fluorescence observed during crop contractions; scale bar: 100 μm. **b**, analisys of dual color images of crops expressing *Subdued*-Gal4>UAS-RGECO. Calcium signal and movement are again significantly correlated confirming the pivotal role of Subdued in peristalsis (n=4; Spearman rho and p are respectively: r=0.52, p=0; r=0.41, p=0; r=0.23, p=0.004; r=0.58, p=0). **c**, plot depicting the frequency of crop contractions in recorded movies (each dot corresponds to a different crop): the frequency of crop peristalsis displayed similar values to the one observed for midgut-related peristalsis (as shown in fig.1D). **d**, snapshots at different timepoints obtained from supplementary video 12 highlight an evident increase in fluorescence coupled to peristalsis in the hindgut, where longitudinal muscles are absent, highlighting again as peristalsis is driven by the *Subdued* activation in circular muscles. Scale bar: 500 μm.

**Supplementary Video 1**

*Drosophila* adult midgut isolated in a mildly acidic solution displays strong non-peristaltic contractions. Scale bar: 700 µm.

**Supplementary Video 2**

Representative movie of an adult fly midgut isolated and recorded in a saline solution at pH=5.5 for 2 min. The Strong whip-like contractions are evident.

**Supplementary Video 3**

Excised guts displayed no superstaltic activity in the first few minutes when exposed to the artificial solution, showing a yellow color in the copper cell region.

**Supplementary Video 4**

The same gut as the one showed in movie S4 displayed peristaltic contractions after ≈10 min exposure to the same solution. The appearance of superstalsis corresponded to a shift in the BPB color in the copper cell region, indicating a perfusion of the artificial solution into the gut lumen.

**Supplementary Video 5**

Monitored midguts expressing the calcium reporter RGECO in all visceral muscles (mef2-Gal4>UAS-RGECO) display longitudinal muscles activation concurrently to superstalsis and circular muscles activation coupled to peristalsis.

**Supplementary Video 6**

Dual color imaging of isolated guts expressing RGECO selectively in longitudinal muscles unveiled the significant coupling of this muscle subsets to superstaltic contractions.

**Supplementary Video 7**

Dual color imaging of isolated guts expressing RGECO under the *Subdued* control displayed significant correlation of peristaltic contractions to increased fluorescence signals. Scale bar: 100 µm.

**Supplementary Video 8**

As in movie S7, is evident a calcium signal increase coupled to peristaltic contraction in *Subdued*-Gal4>UAS-RGCO guts. Scale bar: 100 µm.

**Supplementary Video 9**

Isolated midguts expressing RGECO under Subdued control displayed increased fluorescence upon peristaltic but not superstaltic events, indicating *Subdued* as a specific marker for peristalsis, not responsible for superstalsis.

**Supplementary Video 10**

Isolated crop from adult flies expressing RGECO under the control of *Subdued* also displayed increased fluorescence concurrently to contractions.

**Supplementary Video 11**

Isolated hindguts, where longitudinal muscles are absent, displayed *Subdued*-driven calcium waves coupled to peristaltic contractions. Scale bar: 700 µm.

**Supplementary Video 12**

Movie showing Calcium waves (under *Subdued* drive) spreading throughout the midgut not leading to peristaltic events.

**Supplementary Video 13**

Recording of exposed midguts of flies infected with *Ecc*15 shows both peristaltic and superstaltic events. Scale bar: 500 µm.

**Supplementary Video 14**

Monitoring exposed midguts in starved (24h) flies showed the presence of superstaltic but not peristaltic contractions. Scale bar: 600 µm.

**Supplementary Video 15**

recording of an exposed gut of a fly fed with sugar displayed peristalsis but not superstalsis

**Supplementary Video 16**

Monitoring gut contractions in intact immobilized larva displaying strong superstalsis on top of peristalsis, while the cuticle is not moving. Scale bar: 300 µm.

## Notes

### Competing Interest Statement

The authors have declared no competing interest.

